# Genomic heritability estimates in sweet cherry reveal non-additive genetic variance is relevant for industry-prioritized traits

**DOI:** 10.1101/233296

**Authors:** J. Piaskowski, Craig Hardner, Lichun Cai, Yunyang Zhao, Amy Iezzoni, Cameron Peace

## Abstract

**Background:** Sweet cherry is consumed widely across the world and provides substantial economic benefits in regions where it is grown. While cherry breeding has been conducted in the Pacific Northwest for over half a century, little is known about the genetic architecture of important traits. We used a genome-enabled mixed model to predict the genetic performance of 505 individuals for 32 phenological, disease response and fruit quality traits evaluated in the RosBREED sweet cherry crop data set. Genome-wide predictions were estimated using a repeated measures model for phenotypic data across 3 years, incorporating additive, dominance and epistatic variance components. Genomic relationship matrices were constructed with high-density SNP data and were used to estimate relatedness and account for incomplete replication across years.

**Results:** High broad-sense heritabilities of 0.83, 0.77, and 0.75 were observed for days to maturity, firmness, and fruit weight, respectively. Epistatic variance exceeded 40% of the total genetic variance for maturing timing, firmness and powdery mildew response. Dominance variance was the largest for fruit weight and fruit size at 34% and 27%, respectively. Omission of non-additive sources of genetic variance from the genetic mode resulted in inflation of narrow-sense heritability but minimally influenced prediction accuracy of genetic values in validation. Predicted genetic rankings of individuals from single-year models were inconsistent across years, likely due to incomplete sampling of the population genetic variance.

**Conclusions:** Predicted breeding values and genetic values a measure revealed many high-performing individuals for use as parents and the most promising selections to advance for cultivar release consideration, respectively. This study highlights the importance of using the appropriate genetic model for calculating breeding values to avoid inflation of expected parental contribution to genetic gain. The genomic predictions obtained will enable breeders to efficiently leverage the genetic potential of North American sweet cherry germplasm by identifying high quality individuals more rapidly than with phenotypic data alone.

## BACKGROUND

Sweet cherry (*Prunus avium* L.) is a lucrative fresh market horticultural crop whose monetary worth is directly and indirectly determined by several horticultural and fruit traits. Worldwide, more than 2.8 million tons of sweet cherry fruit were produced in 2014 [1]. In 2015, the U.S. was the second largest producer of cherries, producing 338.6 kilotons of fruit valued at $703 million, of which 60% were grown in Washington State [2,3].

Sweet cherry cultivars must garner a positive critical reception among growers, market intermediaries (a category which includes packers, shippers, and marketers), and consumers to succeed commercially. The U.S. sweet cherry industry and consumers have previously prioritized which fruit trait thresholds are essential for a successful cultivar. Sweet cherry producers identified fruit size, flavor, firmness, and powdery mildew resistance as trait priorities in a survey conducted in 2011 [4]. Powdery mildew (causative agent *Podosphaera clandestine*) is a foliar and fruit disease with a high cost of control in susceptible cultivars. Sweetness and flavor were ranked by consumers as the most important attributes in sweet cherry, followed by firmness, shelf life, and fruit size [5]. Consumers are willing to pay more for sweet, firm cherries with an ideal balance of sweetness and acidity. Sweetness and acidity are quantified with assays for soluble solids content (SSC) and titratable acidity (TA), respectively [5–8]. Market intermediaries indicated a willingness to pay producers more per pound for fruit greater than 2.5 cm in diameter, firmness above 300 g/mm, and SSC above 18° Brix [9]. Market intermediaries also ranked fruit size as the most important trait, followed by firmness and external appearance [10]. The USDA Agriculture Marketing Service evaluates skin color, fruit size, and fruit firmness when grading sweet cherries [11], an assessment which influences market receipts for that crop.

Many of the trait thresholds identified by consumers and the cherry industry alike have been individually met or exceeded through genetic improvement. Beginning with the 1952 release of ‘Rainier’, a highly popular sweet cherry cultivar, the Washington State University sweet cherry program (formerly USDA-ARS) has released several dozen cultivars with improved flavor, size, and firmness in each subsequent release [12,13]. This program and others have largely relied on phenotypic selection complemented with trait-predictive DNA tests for high heritability traits, such as fruit skin color and self-compatibility [13–16]. The Washington State University breeding program has seen genetic gains in fruit dimensions, firmness and other traits of breeding relevance due to moderate heritability of those traits [17–19].

Sweet cherry has a juvenility period of three to five years before a tree is capable of flowering and producing fruit [20]. Therefore, the pace of cultivar release is slow, taking 15 to 25 years between making a cross to cultivar release [16]. Sweet cherry breeding is structured like many other crops: an initial set of crosses is made, followed by evaluation of a large number of offspring. After a rapid screening, the majority of these offspring is discarded, and the remaining selections are evaluated more extensively in replicated trials. Selections are clonally propagated in subsequent evaluations. Consequently, the genetic potential identified in F1 seedlings remains fixed throughout the evaluative phases of a breeding program and is not lost during recombination and segregation.

Understanding the genetic architecture of crop traits can help plant geneticists and allied scientists maximize genetic gain and elucidate the genetic potential of seedlings and parents. Best linear unbiased prediction (BLUP) is an analysis tool that is used to estimate the genetic potential of each individual from unbalanced trials by modeling genetic effects as a random effect in a mixed model [21]. It requires prior estimation of genetic variance components, which are obtained through maximum likelihood, restricted maximum likelihood (REML) or Bayesian approaches [22,23]. Pedigree-based BLUPs have been developed to leverage information from related individuals. This is used to estimate the genetic potential that a parent can pass to its offspring and is termed “breeding value” [24]. Genomic BLUPs (GBLUPs) are an extension of pedigree-based BLUPS, using DNA marker information instead of pedigree information to construct a realized relationship matrix between individuals in a population. The realized relationship matrix can more accurately estimate relatedness, particularly among full siblings, than the pedigree-based relationship matrix [25–27]. The resultant breeding values are expected to more closely mirror the true genetic potentials of individuals [28–30].

Breeding values derived from BLUPs have been used to successfully identify superior individuals in several rosaceous crops including apple, peach, raspberry, and strawberry [31–37]. Extensive work has been done in apple to estimate the breeding values from unreplicated trials [31,33,38,39]. Breeders have observed enhanced genetic gain using both pedigree-based and genome estimated breeding values in other perennial tree crops, including citrus, rubber and *Eucalyptus* [40–43]. Sweet cherry shares many of the breeding scheme challenges of apple and other perennial tree crops: unbalanced trials and a long juvenility period. Hence, the same methodologies can be utilized.

Additive effects are considered to be the largest component of genetic variance that is passed to progeny [44]. While many genome-wide approaches including GBLUPs have been employed to estimate breeding values across crops, these methods are almost solely focused on estimating additive effects alone as a proxy for total genetic effects. Few studies have examined non-additive genetic variance components in rosaceous crops [45]. Kumar et al. [45] reported on a comprehensive study estimating sources of genetic variance for 32 traits in apple across 17 families and two locations using GBLUPs.

In cherry, there are few published accounts that utilize BLUPs or other genome-wide DNA-enabled approaches for estimating the genotypic value of individuals. The only published genome-wide study in sweet cherry estimated breeding values for cherry fruit size in U.S.-relevant germplasm from large-effect QTLs in a Bayesian analysis, but it did not include genetic background effects [18]. There is no published information on the genome-wide additive and non-additive variance components and prediction of the genetic value of individuals for any sweet cherry trait.

This study addresses a deficiency of published information on genetic parameters for sweet cherry breeding-relevant traits beyond those influenced primarily by large-effect QTLs by obtaining robust estimates of genetic variance components. To ensure wide applicability of the study for cherry, we used a large set of sweet cherry breeding germplasm. These data were gathered from germplasm in public sweet cherry breeding programs as part of RosBREED project [46]. Our objectives were to: (1) estimate variance components across a broad spectrum of traits in sweet cherry germplasm important to North American breeders and producers, and (2) assess the predictive accuracy of obtained genome-estimated breeding values (GEBVs) for a subset of the most valuable traits. Previous studies show that the genome-estimated breeding values of individuals that are robust across years and families can increase the pace and efficiency of breeding. Specifically, valuable cherry parents can be identified more quickly and with greater confidence than those obtained through phenotypic data alone.

## METHODS

### Germplasm

We used all individuals from the RosBREED sweet cherry Crop Reference Set with genome-wide SNP data, totaling 505 individuals (Additional File 1). This set consisted of cultivars (n = 42), wild accessions (n = 3), unreleased selections (n = 24), and unselected offspring (n = 436) from 66 families. The unselected offspring category includes 77 F1 offspring derived from a wild parent and 359 F1 offspring derived from existing cultivars. Trees were grown at two sites in Washington State (U.S.A.) located approximately 0.5 km apart: the Irrigated Agriculture Research and Extension Center of Washington State University Roza Unit, (46 □29’N and 119 □73’W) and at Pear Acres (46 □29’N and 119 □75’W). Each tree was planted in 2006, 2007, or 2008 and managed using conventional orchard management practices. Unselected offspring were grown on their own roots, and the remaining germplasm were grown on Gisela 6 rootstock [47]. A single tree was used for each individual. The Crop Reference Set was established to represent North American sweet cherry breeding germplasm for QTL identification and validation and other quantitative genetics endeavors [48].

### Phenotypic data

This study used the sweet cherry phenotypic data set previously described in Chavoshi et al. [49] obtained in the RosBREED project. This data set consisted of 32 traits evaluated in 2010, 2011, and 2012. Standardized phenotyping protocols for sweet cherry [49] were used. For individual fruit traits, the five largest fruit without blemish were measured and averaged. In the case of pitting and cracking, the proportion of fruit observed with symptoms out of 25 fruit was recorded. Bulked fruit traits (bulked fruit weight, bulked firmness, bulked SSC, and bulked TA) were reported as the average of measurements over 25 fruit.

Nine traits of the 32 were focused on here because of their importance in new sweet cherry cultivars: time to bloom, time to maturity, pedicel-fruit retention force (PFRF), fruit dimensions, fruit weight, firmness, SSC, TA, and powdery mildew incidence. Time to bloom and time to maturity were measured both in Julian calendar days starting from January 1^st^ of the calendar year and in growing degree days (GDD). The force required to pull a ripe cherry fruit from its pedicel, PFRF, and fruit weight were both measured in grams. Firmness, SSC, and TA were measured in units of g/mm, Brix°, and percentage, respectively. Foliar powdery mildew incidence was scored in August of each year, immediately after the fruiting season, on a 0-5 scale, where 0 is no infection and 5 is highly infected leaves. These nine traits are referred to as “focus traits” for the rest of the study. All trait data were measured over three years except for powdery mildew incidence, which was not assayed in 2010. Results from the other traits are given in the supplementary material, but not discussed.

Several transformations of the trait data were performed for the focus traits. “Fruit dimensions” was determined newly here as the first component from a principal component analysis between fruit length and fruit width, which are both end-to-end fruit measurements in millimeters. The first principal component summarized 95.4% of total phenotypic variation for fruit length and width. Growing degree days was calculated for an alternative measure of phenological traits. Climatic data was obtained from Washington State University’s AgWeatherNet using the “Roza” station [50], using a base temperature of 4.5 °C and maximum of 30 °C. Daily maximum temperatures above 30 °C were reduced to 30 °C, and negative temperatures were set to zero, following McMaster and Wilhelm [51]. Erroneous data points, defined as those larger than twice the next largest value or less than one-half of the next smallest value and having a studentized residual with an absolute value greater than 5, were removed. Such data were assumed to be data entry errors. There were 97 individuals with no phenotypic data: 13 selections and 84 unselected progeny. These individuals were used in the model-building and prediction steps for all models except for cross validation.

### SNP data

The SNP data were obtained from the RosBREED project using the RosBREED cherry 6K SNP array v1 (an Illumina Infinium^®^ II array) [52]. The SNP curation pipeline is described in Cai et al [53]. Missing data were imputed with Beagle as implemented in SynBreed [54,55] using the hidden Markov model and a minor allele frequency of 0.05. Individuals or SNPs missing more than 25% data were removed from analysis and the SNP. In total, a genome-wide set of 1615 SNPs was used.

### Statistical modeling

Variance components were estimated with R-ASReml 3.0 [56], and additional statistical analyses were conducted in R v3.4 [57]. The following model was used for initial estimates of genetic effects for a single trait, **Y**:

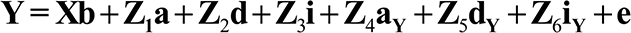

where **a, d, i, a_Y_, d_Y_** and **i_Y_** are the random variables for additive effects, dominance effects, effects from additive-by-additive epistatic, additive-by-year effects, dominance-by-year effects, and epistasis-by-year effects, respectively. Variables **Z_1-3_** and **Z_4-6_** are design matrices for main effects and interaction terms, respectively. Dimensions of **Z_1-3_** are *nY×Y* and **Z_4-6_** are *nY×nY*, where *n* is the number of individuals and *Y* is the number of years with trait data for an individual. Year was treated as a fixed effect, where **X** is the design matrix relating observations to years and b is a vector of fixed effects due to year. In a preliminary analysis, the effect of location was evaluated as a fixed effect using a Wald test. Location did not have a significant effect on the focus traits (p-value > 0.10) and was omitted from the model. Random variables were assumed to follow a normal distribution:

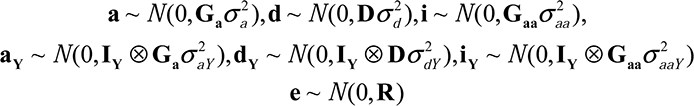

The covariance structure for year was modeled as a repeated measure: **R** = **I**_*individual*_ ⊗ **e_Y_** where **I**_*individual*_ is an identity matrix of individuals included in the study and **e_Y_** is a 3×3 matrix of year error terms using a general correlation structure implemented in ASReml. The genomic additive relationship matrix was computed with R/rrBLUP [58] using the VanRaden method [59]:

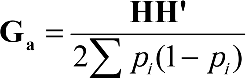

where p_i_ is frequency of the positive allele for a single marker column, and **H** was computed as equal to centered marker data, {*H*} _*ij*_= {*M*}_*ij*_ − 2(*p_i_* − 0.5). **M** is an *n* x *m* marker matrix with *n* individuals and *m* markers expressed as (-1,0,1) frequency. The dominance relationship matrix was computed using normalized matrices described by Su et al. [60] and implemented using a custom R program [61]:

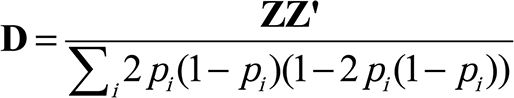

where the **Z** matrix is a transformation of the marker matrix, **M**:

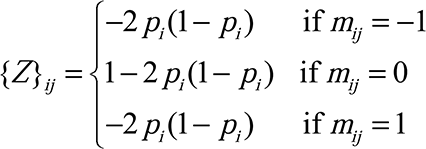

The epistatic relationship matrix for additive by additive effects was computed by taking the Hadamard product between **G_a_**, the additive genomic relationship matrix, and itself: **G_aa_** = **G_a_** °**G_a_**.

When a relationship matrix was not positive definite, a small constant of 1e^−6^ was added to the first eigenvector, and the matrix was inverted.

The full model included additive, dominance, and epistatic main effects and their interactions with year and is also called the “ADI model” in this paper. Model fit was assessed by checking for model convergence, examining studentized residuals for each trait by year combination, and examining the extended hat matrix for influential observations. The default model convergence criteria for ASReml were used, in which the final iteration must satisfy the following conditions: a change log likelihood less than 0.002 * previous log likelihood, and the variance parameters estimates change less than 1% from the previous iteration. The extended hat matrix for linear mixed models is:

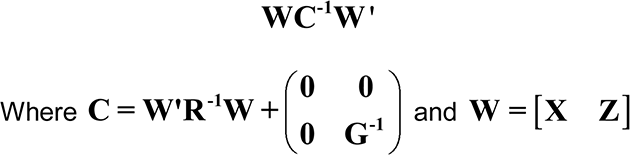

Influential data points were those with a value greater than 2 times the average value of the diagonal of the hat matrix excluding zeros.

The statistical significance of main effects and interactions were tested by first generating reduced models and then performing log-likelihood ratio tests between full and reduced models. To account for positively-bound variance component estimates, a mixture of Chi-square distributions as implemented in the R package asremlPlus [62] was used. Non-significant values from the log likelihood ratio tests were interpreted as the reduced models being as effective as the full model in modeling the response variable. Heritability numerators were estimated as 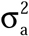 for narrow-sense heritability (*h*^2^) and as 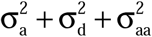 for broad-sense heritability (*H*^2^); both were divided by the sum of the variance components for final heritability estimates. Genetic values were computed as the sum of main effects for **a, d** and **i** for an individual, following the methodology of Kumar et al. [45]. Genotype-by-year effects are the sum of **a_y_**, **d_y_**, and **i_y_** when all years were used in the estimation.

### Model validation

Five-fold cross validation was used where the data set was randomly divided into 5 equal-sized parts (“folds”), a single fold (20% of the individuals) was removed across all years, and the remaining observations were used for variance component estimation and prediction of genetic values. The resultant model was used to predict genetic values of those removed individuals. This process was repeated for all 5 folds. Observations lacking phenotypic information for a specific year and trait were excluded from the model-building and validation. Because predictions can be affected by sampling variance, 5-fold cross validation was repeated 25 times using different randomly generated folds for each iteration. In addition, cross validation was performed, omitting each of the 66 full-sib families or a year as validation populations. These latter situations were intended to reflect the situation of predicting genetic performance for previously unphenotyped individuals that are related to the training population, and for predicting performance for an unobserved year. Prediction accuracy was assessed by computing correlation coefficients between predicted genetic values and observed data adjusted for fixed effects.

### Other statistics

The statistical significance of year on the models was checked with the Wald test. Genetic-by-year effects were further explored by estimating genetic values and genetic variance components using a single year of data. Spearman’s rank-order correlations were conducted to evaluate changes in rank of genetic values of individuals across years. Pairwise Pearson (r) and Spearman (ρ) correlations between traits were assessed for the multi-year ADI model. Principal component analyses were conducted on correlation matrix of genetic values calculated from (1) all individuals used in this study, and (2) only the cultivars and ancestors (n=48), using 8 independent traits: bloom time, harvest time, pedicel-fruit retention force, fruit weight, firmness, SSC, TA, and powdery mildew incidence. The first and second principal components were graphed on a biplot [63], where the rotations for plotting the variables were scaled by the first eigenvalue.

## RESULTS

### Distribution of phenotypic data

All trait distributions (consisting of 600-755 data points for each trait) were influenced by the year of data collection (Fig. 1). Wald test results for year were consistently highly significant for all focus traits across all models (p < 0.001 in all cases).

**Figure 1:**
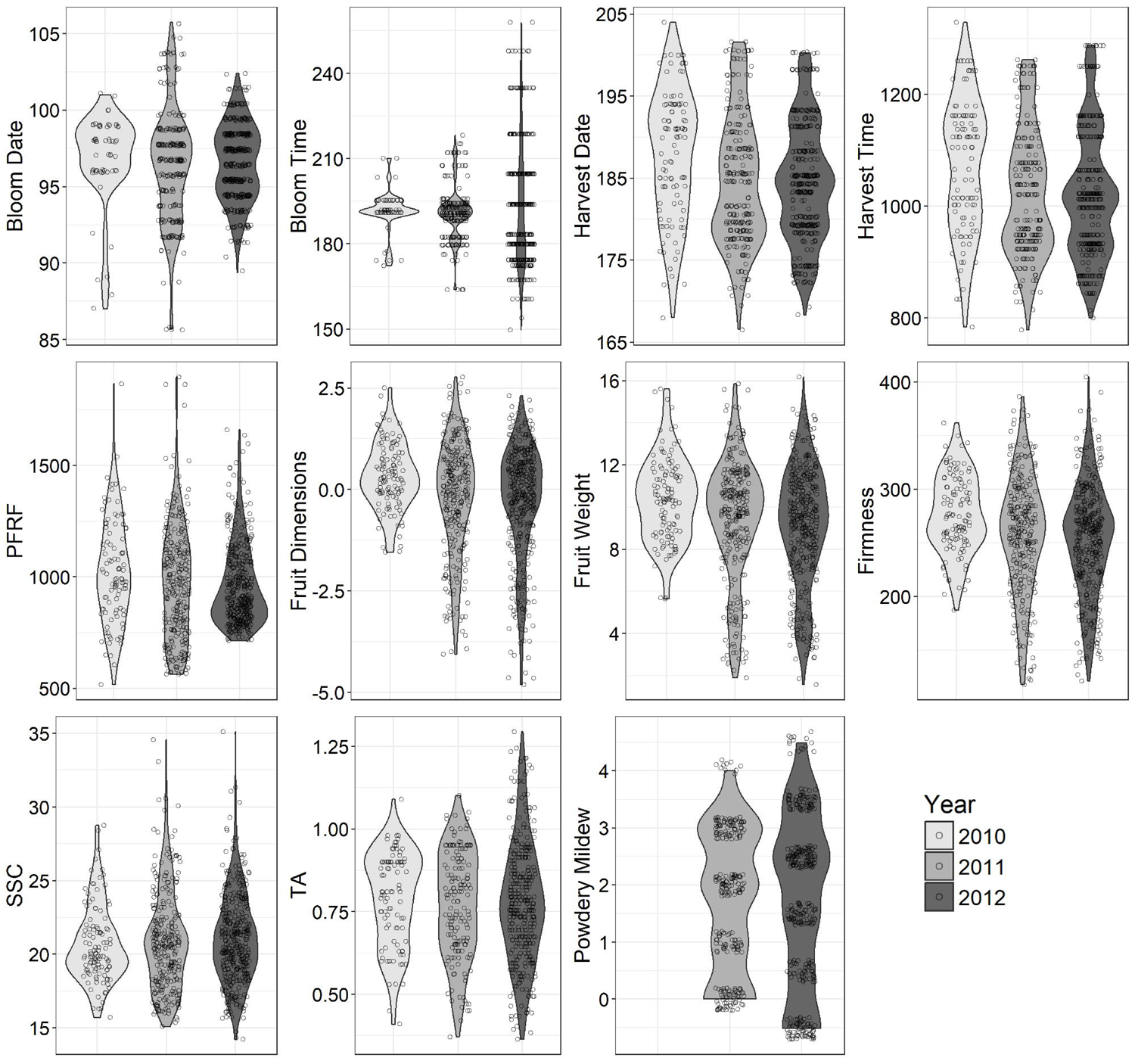
Violin plots of nine traits by year, adjusted for fixed effects due to year and overlaid with the observations from each year.

The 2010 data visually differed most from the other years, particularly for bloom date, fruit dimensions, fruit weight, firmness, and SSC. Data in 2010 were also the most sparse compared to data from other years (Additional File 2). Fruit dimensions and fruit weight had similar distributions across years. Although the distributions of bloom date and bloom time seemed to differ, the accumulation of GDD remained relatively stable over the three years. However, GDD accumulation was higher in early 2010 than other years during the critical period of flower bloom (data not shown).

### Statistical assumptions and model fit

All models for the focus traits converged. Inspection of the residual plots and quantile-quantile plots signal that the error terms were independently and identically distributed (results not shown). The extended hat matrix revealed no influential data points for any of the models. Appropriate residual patterns were observed for all models and traits (results not shown), demonstrating no major departures from the assumption of homoscedasticity. Moderate correlations were observed between the additive, dominance and epistatic effects within a trait for the full model (r = 0.3 − 0.7). Population structure was observed among the individuals. In a principal component analysis of the correlation matrix of the SNP data, the first two components summarized 14% of the variation. There was distinct grouping of the wild accessions and offspring derived from those wild accessions along the second principal component (data not shown). Visual inspection of the diagonals and off-diagonals from the realized relationship implies a single Gaussian distribution of the matrix elements. Thus, the population structure had minimal impact on the genomic additive relationship matrix (Additional File 3).

Log likelihood ratio tests comparing reduced models with the full ADI model demonstrated that the full model was not necessary to describe trait variance for any focus trait (Table 1). The main effects-only model that included only additive, dominance, and epistatic effects was significantly different from the full model (p-values <0.05) for all focus traits, except for powdery mildew incidence and SSC, which had notable p-values defined as less than 0.10. Reduced models consisting of single main effects (additive, dominance or epistatic) or single main effects plus their year interaction term (e.g., additive and additive-by-year) were highly significant for all traits. This demonstrates that the reduced models did not adequately capture variation compared to the full model. For most focus traits, genetic models that included additive, epistatic, additive-by-year and epistasis-by-year effects were not statistically different from the full model. Thus, dominance and dominance-by-year could be dropped from their genetic models without significant loss of information. Traits that were exceptions to the above were fruit weight, fruit dimensions, and bloom date, for which optimal fit was obtained by including dominance in the model. For all traits, dominance-by-year and epistasis-by-year effects could be removed from the model without much loss of information. Additive-by-year effects had a statistically significant effect on bloom date, bloom time, and PFRF (p<0.01).

**Table 1:**
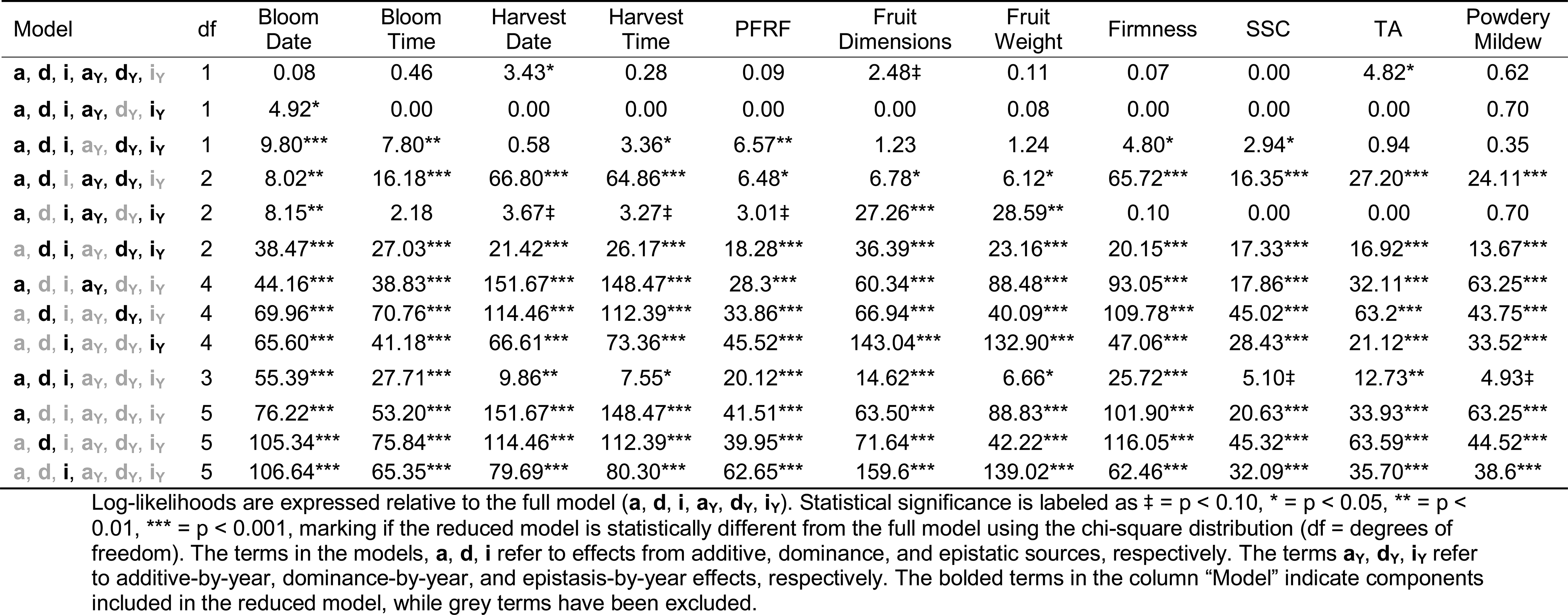
Log-Likelihood ratio test statistics for reduced models.

### Genetic variance and predictive ability of full model

Variance component estimates from the full model indicated moderate to high broad-sense heritabilities across the focus traits, ranging from 0.47 for pedicel-fruit retention force to 0.83 for harvest date (Table 2). Narrow-sense heritabilities ranged from 0.20 for PFRF to 0.37 for fruit dimensions. Epistasis was the single largest genetic variance component for most traits: bloom time (28%), harvest date (48%), harvest time (48%), firmness (49%), SSC (27%), TA (33%), and powdery mildew incidence (42%). Additive variance was the largest component for bloom date (37%), PFRF (20%), and fruit dimensions (37%). Dominance was the largest variance component only for fruit weight (34%); in contrast, dominance represented less than 1% of trait variance for firmness, SSC, TA, and powdery mildew incidence. Genotype-by-year effects were less than 10% for all traits except bloom date (**a_Y_** = 11%) and TA (**i_Y_** = 14%). Residual variance of most traits was less than 25% of phenotypic variance, except for PFRF (45%) and SSC (48%). Variances and standard errors for all components and traits, and variance percentages, are provided in Additional Files 4 and 5, respectively.

**Table 2:** Variance components (%), narrow-sense heritability (*h*^2^), broad-sense heritability (*H*^2^), the coefficient of correlation (*r*), the coefficient of correlation after cross validation (*r_cv_*), and the total number of observations for model building (N).

|  | Bloom<br>Date | Bloom<br>Time | Harvest<br>Date | Harvest<br>Time | PFRF | Fruit<br>Dimensions | Fruit<br>Weight | Firmness | SSC | TA | Powdery<br>Mildew |
| --- | --- | --- | --- | --- | --- | --- | --- | --- | --- | --- | --- |
| variance component (%) |  |  |  |  |  |  |  |  |  |  |  |
| additive (A) | 33.20 | 25.45 | 27.39 | 27.87 | 19.83 | 37.40 | 30.76 | 27.49 | 21.59 | 27.19 | 28.31 |
| dominance (D) | 10.80 | 11.48 | 7.73 | 6.68 | 11.10 | 26.80 | 33.61 | 0.42 | 0.00 | 0.00 | 0.00 |
| epistasis (I) | 17.47 | 27.84 | 47.66 | 47.90 | 15.62 | 8.36 | 12.08 | 48.96 | 26.76 | 32.81 | 41.52 |
| A x Year | 11.16 | 8.10 | 1.18 | 2.89 | 6.30 | 2.98 | 2.26 | 4.57 | 4.08 | 3.42 | 1.57 |
| D x Year | 4.23 | 0.00 | 0.00 | 0.00 | 0.00 | 0.00 | 0.52 | 0.00 | 0.00 | 0.00 | 1.19 |
| I x Year | 2.06 | 3.63 | 5.71 | 1.65 | 2.31 | 8.03 | 1.14 | 0.94 | 0.00 | 14.15 | 4.33 |
| error | 21.08 | 23.48 | 10.33 | 13.02 | 44.84 | 16.42 | 19.62 | 17.62 | 47.56 | 22.43 | 23.07 |
| trait heritability and genome-estimated breeding values accuracy |  |  |  |  |  |  |  |  |  |  |  |
| $h^2$ | 0.33 | 0.25 | 0.27 | 0.28 | 0.20 | 0.37 | 0.31 | 0.27 | 0.22 | 0.27 | 0.28 |
| $H^2$ | 0.61 | 0.65 | 0.83 | 0.82 | 0.47 | 0.73 | 0.76 | 0.77 | 0.48 | 0.60 | 0.70 |
| $r$ | 0.88 | 0.90 | 0.97 | 0.97 | 0.83 | 0.94 | 0.95 | 0.94 | 0.82 | 0.88 | 0.93 |
| $r_{CV}$, 5-fold | 0.56 | 0.48 | 0.78 | 0.79 | 0.59 | 0.78 | 0.77 | 0.69 | 0.46 | 0.42 | 0.68 |
| $r_{CV}$, -year | 0.58 | 0.48 | 0.88 | 0.88 | 0.58 | 0.82 | 0.83 | 0.76 | 0.47 | 0.50 | 0.74 |
| $r_{CV}$, -family | 0.55 | 0.46 | 0.74 | 0.74 | 0.55 | 0.76 | 0.70 | 0.66 | 0.38 | 0.31 | 0.58 |
| N | 644 | 644 | 665 | 665 | 759 | 774 | 764 | 763 | 768 | 577 | 604 |

Correlations between adjusted phenotypic data and genetic values from the ADI model were high, 0.82-0.97 for all focus traits (Table 2). Coefficients of correlation under cross validation were very similar for 5-fold cross validation and when a year was left out. Correlations for cross validation that omitted full-sib families were the lowest among the three cross validation scenarios. Across all cross-validation scenarios, those traits with the highest broad-sense heritabilities, fruit dimensions, fruit weight, firmness, harvest date, and harvest time, had the most consistently high prediction accuracies (*r* > 0.65). The lowest prediction accuracies were observed for SSC and TA, which never exceeded 0.50.

### Heritability and predictive ability of reduced models

Broad-sense heritability was largely unchanged across the reduced models (ADI to AI and AD) for all focus traits (Fig. 2). Narrow-sense heritability gradually increased with decreasing model complexity for all focus traits, from the full model to the AD model and from the AD to the A model. Narrow-sense heritability was highly similar in the AI and ADI models for all traits except for fruit dimensions and fruit weight, in which the AI *h^2^* was noticeably higher in the AI model compared to the ADI and AD models (Fig. 2). In the additive effects-only model (A), *H^2^* was similar in value to the *h^2^* of the other models.

**Figure 2:**
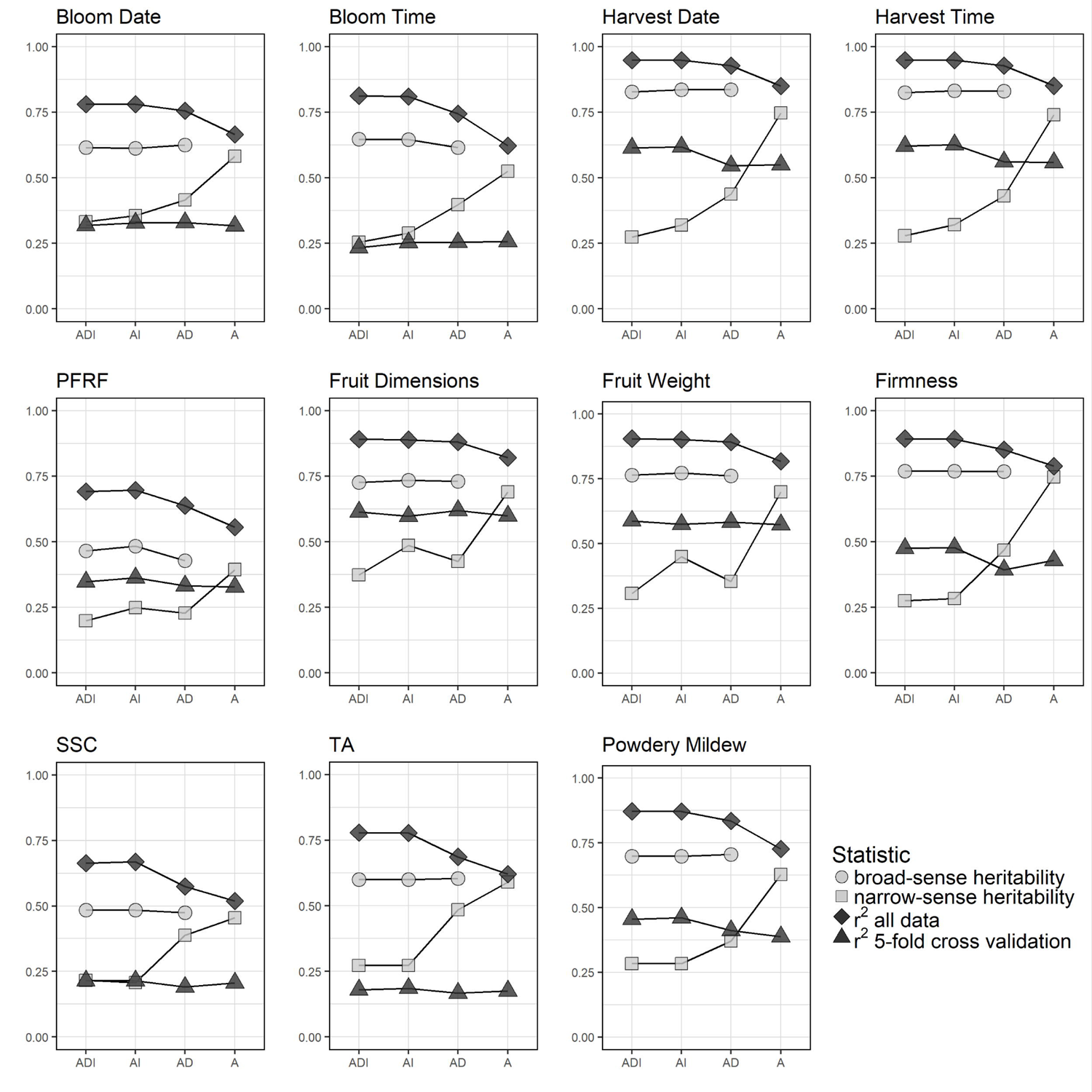
Heritability and predictive ability of four genetic models for each of nine focus traits.

Predictive power, as measured by *r^2^*, was consistent between the ADI model and the AI model for all traits (Fig. 2). The predictive power decreased slightly for the AD model compared to the full model, and decreased slightly more for the A model compared to the AD model. The *r^2^* values under 5-fold cross validation varied little across genetic models for all traits, only decreasing slightly in the AD and A reduced models for harvest date, harvest time, and firmness. Spearman rank correlations between the full and reduced models indicated minimal changes in rankings of individuals when using the AD and AI models (r = 0.96-1.00) and small changes in the A model compared to ADI model (r = 0.91–0.96) for genetic values and breeding values (Additional File 2).

### Single year analysis

Variance components estimated with a single year of data varied substantially across years for all focus traits (Fig. 3). For all traits except harvest date and harvest time, the percentages of additive variance differed by 10% or more across years. Additive variance for harvest date and harvest time varied the least among the focus traits, 37 to 44% and 37 to 47%, respectively. Dominance variance components for SSC and TA were close to zero (<0.0001%) across all years, while at the other extreme, dominance variation for fruit dimensions was always greater than 20%. Epistatic variance consistently composed a large percentage of genetic variance for firmness (>32%) and powdery mildew incidence (>49%). Genotype-by-year effects were greatest for TA (18%), bloom date (18%), and bloom time (12%).

**Figure 3:**
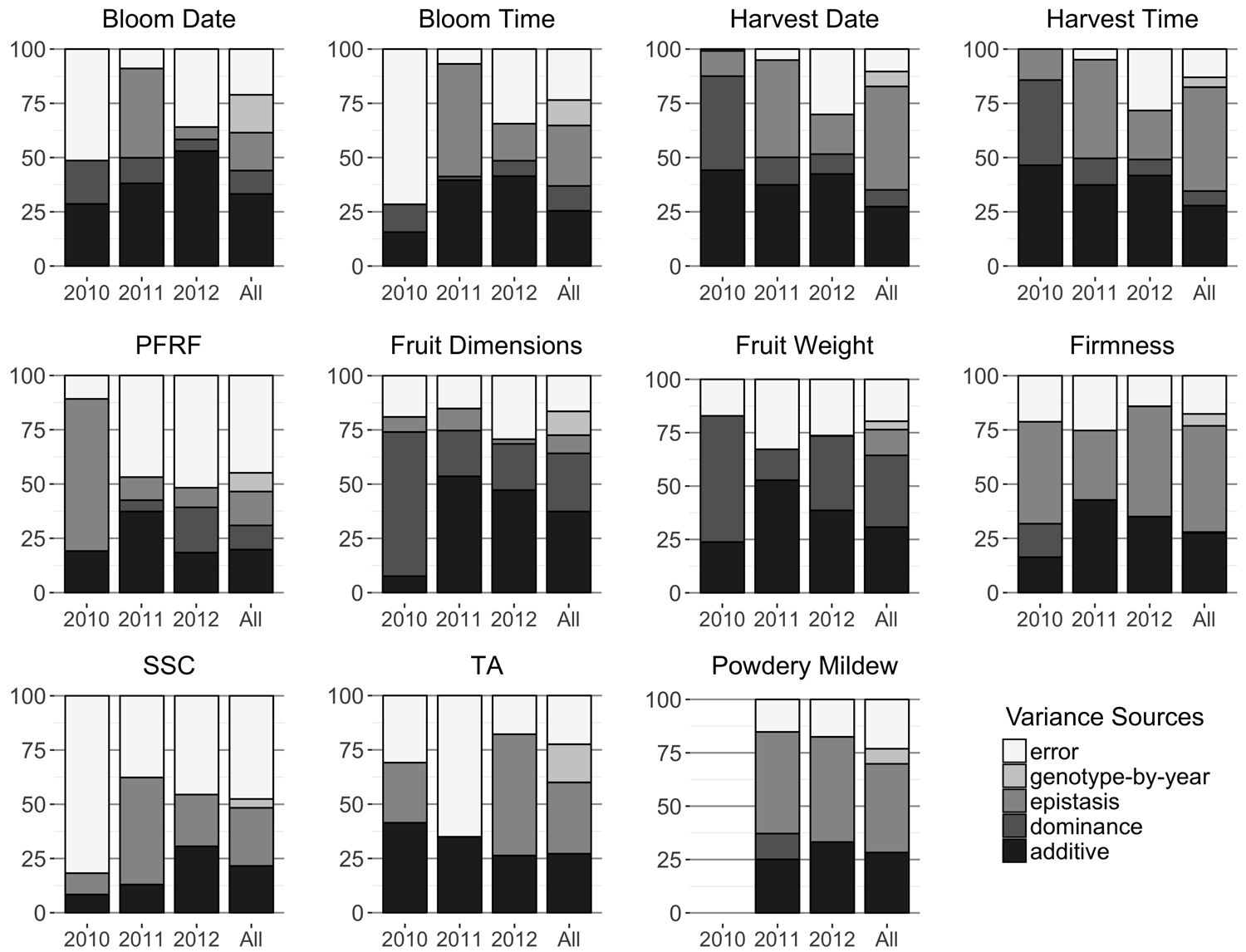
Variance components by each year and across years for the full (ADI) model.

Rankings of individuals by genetic values estimated from each a single year of data significantly differed from the multi-year genetic rankings in Spearman rank correlation tests (p <0.001, Additional File 2). Rank correlations between the 2010-derived predictions and the multi-year predictions were lower than the subsequent years (2010: 0.35–0.63; 2011: 0.58–0.92; 2012: 0.85–0.97). However, correlations between single-year breeding values and phenotype implied a better fit for all years and traits than the single-year breeding values with their multi-year counterparts (ρ = 0.64-1.00) (Additional File 2).

### Correlations among trait genetic values

The genetic values of the focus traits had weak to moderate positive correlations with each when considering only unreleased offspring and selection, with some exceptions (Table 3). Fruit weight and fruit dimensions, harvest date and harvest time, and bloom date and bloom time were all highly correlated pairs of traits (r > 0.90, Table 3). SSC was negatively correlated with all focus traits except TA. Titratable acidity was also negatively correlated with fruit dimensions, fruit weight and powdery mildew incidence. In a biplot of the correlation matrix of the named cultivars using eight independent traits, the first two principal components summarized 55% of the variance (Fig. 4). All variables but SSC and TA skewed to the left, corresponding to the negative correlations between SSC and all variables except TA. Wild ancestors and wild offspring were on the right side of the biplot corresponding to their high SSC, low powdery mildew incidence, and low fruit weight. Additional figure 6 further separates the sweet cherry founders and derived cultivars by fruit weight and SSC content.

**Table 3:** Pairwise trait correlations and covariances between genetic values for sweet cherry selections and unselected offspring.

|  | Bloom Date | Bloom Time | Harvest Date | Harvest Time | PFRF | Fruit Dimensions | Fruit Weight | Firmness | SSC | TA | Powdery Mildew |
| --- | --- | --- | --- | --- | --- | --- | --- | --- | --- | --- | --- |
| Bloom Date | **3.507** | 0.897*** | 0.317*** | 0.314*** | 0.301*** | 0.136* | 0.196*** | 0.223*** | -0.101 | 0.133* | 0.184** |
| Bloom Time | 18.17 | **117.1** | 0.213*** | 0.208*** | 0.198*** | 0.087 | 0.130* | 0.134‡ | -0.071 | 0.064 | 0.218*** |
| Harvest Date | 3.446 | 13.38 | **33.72** | 0.998*** | 0.255*** | 0.340*** | 0.346*** | 0.547*** | -0.364*** | 0.107 | 0.220*** |
| Harvest Time | 50.93 | 195.4 | 502.2 | **7508** | 0.251*** | 0.334*** | 0.341*** | 0.549*** | -0.355*** | 0.106 | 0.225*** |
| PFRF | 74.88 | 283.9 | 196.2 | 2883 | **17630** | 0.566*** | 0.603*** | 0.465*** | -0.161*** | 0.173‡ | 0.185*** |
| Fruit Dimensions | 0.3082 | 1.142 | 2.394 | 35.10 | 91.09 | **1.468** | 0.946*** | 0.511*** | -0.507*** | -0.210*** | 0.462*** |
| Fruit Weight | 0.9209 | 3.695 | 5.042 | 74.33 | 201.1 | 2.880 | **6.311** | 0.514*** | -0.435*** | -0.208*** | 0.505*** |
| Firmness | 17.26 | 59.91 | 131.1 | 1964 | 2546 | 25.55 | 53.26 | **1702** | -0.392*** | 0.065 | 0.387*** |
| SSC | -0.3403 | -1.387 | -3.793 | -55.10 | -38.44 | -1.101 | -1.958 | -28.99 | **3.214** | 0.267*** | -0.340*** |
| TA | 0.02486 | 0.06856 | 0.06204 | 0.9132 | 2.295 | -0.02541 | -0.05219 | 0.2668 | 0.04778 | **0.009934** | -0.215** |
| Powdery Mildew | 0.3509 | 2.409 | 1.304 | 19.91 | 25.11 | 0.5707 | 1.295 | 16.28 | -0.6215 | -0.02181 | **1.041** |

Correlations and covariances are given in the upper triangle and lower triangle, respectively, and trait variances are bolded on the diagonal.
Statistical significance is labeled as ‡ = $p < 0.10$ , \* = $p < 0.05$ , \*\* = $p < 0.01$ , \*\*\* = $p < 0.001$ , signaling if the correlations are different from zero.

**Figure 4:**
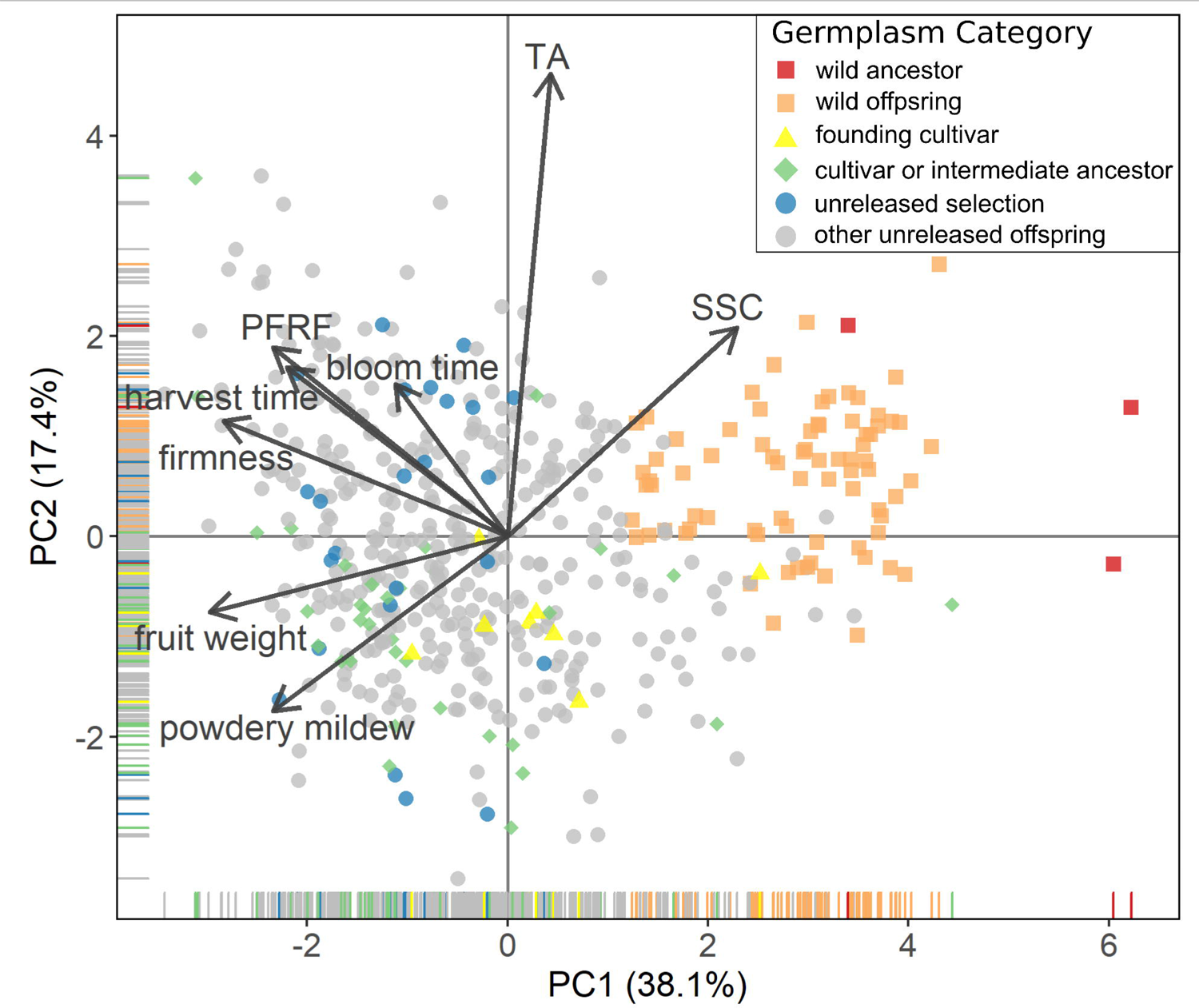
Biplot of genetic values among the RosBREED sweet cherry Crop Reference Set using the correlation matrix of eight traits. Trait rotations were scaled by the first eigenvalue.

## DISCUSSION

Results indicated high broad-sense heritability for all of the focus traits and also illuminated the importance of non-additive variation in the sweet cherry traits studied. A poorly-fitting genetic prediction model can mispresent the genetic variances of traits and the potential for genetic gain.

### Importance of model fit and consequences for predictive ability

This study demonstrated that for most traits, non-additive sources of variation comprised an equal or larger portion of the genetic variance than additive variance. A genetic model including additive, epistatic, additive-by-year and epistasis-by-year effects was usually the most parsimonious approach for capturing major sources of variation. Exceptions were fruit dimensions and fruit weight, which instead were best described by a model with additive, dominance and additive-by-year effects, and harvest date, best described by a main effects-only model.

Using an incorrect model to determine genome-wide breeding values can provide misleading information for making breeding decisions. Table 4 illustrates the consequences of using a poorly-fitting reduced model for estimating breeding values. Breeding values were often larger in relative magnitude in the reduced models compared to the full model, which can exaggerate genetic gains possible in the population. For example, days to maturity in an Ambrunes/Sweetheart cross would be overestimated by twice as many days in the additive-only model compared to the ADI model. Likewise, crosses with the wild accession MIM 23 were predicted to result in midparent values of fruit size twice as small in the A model compared to the ADI model (Table 4). The inflation of additive variance when non-additive sources are omitted has been documented in several other species including apple, loblolly pine, white spruce cassava, cattle, pigs, Coho salmon, and rainbow trout [27,45,60,64–68].

**Table 4:**
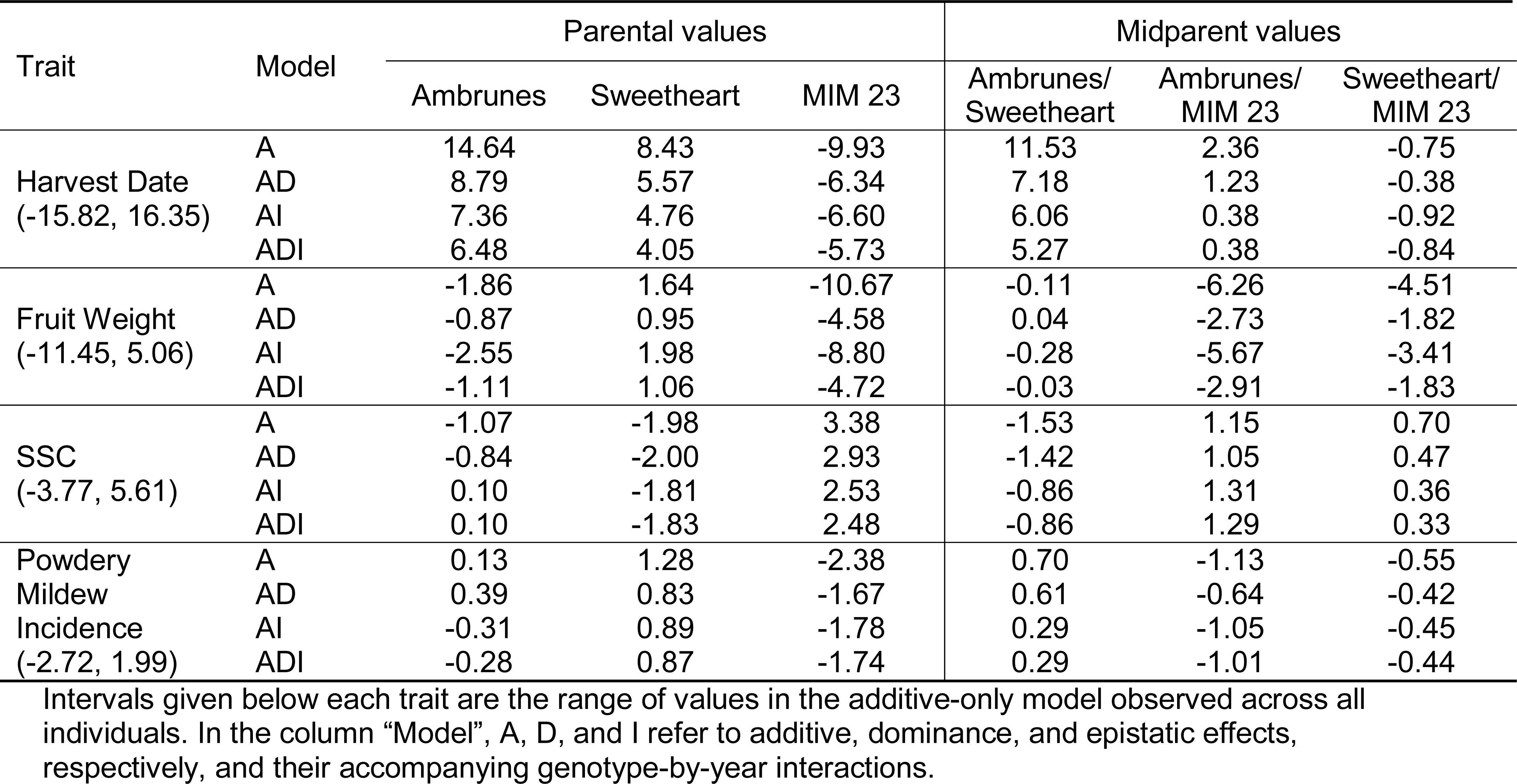
Breeding values and midparent values under different genetic models demonstrated with several individuals and traits.

If genetic values are used to select individuals to be clonally propagated for further trialing or cultivar release, then the genetic model has a lower, perhaps negligible, influence on prediction of total genetic performance. Ceballos et al [69] argued that using total genetic values from additive and non-additive variance components provides greater potential for genetic gain under clonal selection. However, our results showed that the estimated broad-sense heritability and the genetic values of sweet cherry individuals are largely unchanged across the different genetic models. This demonstrates that there is effectively no change in genetic gain if a more complex model is used for identifying high-performing individuals (Figure 2, Additional File 2).

Including year as a main effect was warranted in this study, given the statistically significant effect of year on all traits. However, the effect of including genotype-by-year interactions varied by the trait and genetic variance component. Genotype-by-year interactions were generally of much smaller magnitude than the main genetics effects and largely absent for dominance effects (Table 1, Fig. 3). Nevertheless, year had a major effect on genetic effects estimates and was included as a fixed variable to obtain robust predictions across years. Year often has a statistically significant effect on the traits of sweet cherry and other rosaceous crops, including sweet cherry pedicel-fruit retention force [70], apple fruit texture [71], sugar content in peach and nectarine [72], and several phenological and fruit quality traits in strawberry [73].

This study demonstrated the need for a training population to fully capture variation of the target population in order to maximize prediction accuracy. The single year analysis showed that although a model built using a single year of data could be used accurately to predict individuals evaluated in that year, it could not be easily extrapolated to individuals whose genetic values lie outside the distribution of the training data (Table 2, Additional File 2). The GBLUP approach relies on information from relatives to improve the accuracy of the estimates [74]. Because there were often sparse observations for a single year, sampling error biased the single-year estimates and resulted in models that fit the data within each year, but not across years. These effects were likely exacerbated with wild accession, distantly related cultivars and derivatives from both groups. However, the true pairwise genetic covariance between the distantly related germplasm is estimated with less reliability with the realized relationship matrix than more closely related germplasm [75].

### Genetic architecture of focus traits in sweet cherry

This study confirmed the extensive opportunity in North American sweet cherry germplasm for genetic improvement of the phenological traits of harvest timing and, to lesser extent, bloom timing. Previous QTL studies for fruit maturity date across several *Prunus* species determined bloom timing and harvest timing to be highly heritable with a large-effect QTL on LG4 [76]. Our findings also demonstrate the large broad-sense heritability for these traits – reaching a ceiling of 0.83 for harvest time and 0.65 for bloom date (Fig. 2). There appears to be little advantage to using GDD to Julian days, since pairs of phenological traits for bloom and harvest timing displayed highly similar genetic architecture and predictive accuracy. The data were all gathered from a single location, in which GDD did not vary dramatically during the years of evaluation. This may explain why GDD did not improve the model predictive ability over Julian days (Fig 2, Table 2). Bloom timing has become increasingly important as a trait relevant to productivity, since variable climatic patterns in temperate regions can result in earlier flowering and an increased risk of floral freeze damage [77]. Furthermore, since sweet cherries are a fresh market product that is subject to rapid postharvest deterioration, it is crucial to for sweet cherry breeders and producers to understand the expected time frame for fruit maturation [76]. These results may help sweet cherry breeders identify the best parents in order to target a harvest timing window.

Moderate prospects were observed for genetic improvement of pedicel-fruit retention force (*h*^2^ = 0. 20, *H*^2^ = 0.46, Table 2), where a low PFRF value is sought for mechanical harvest systems. Positive correlations observed between PFRF and fruit dimensions, fruit weight, and firmness (Table 3) contrasted with findings by Zhao et al. [70], in which PFRF was largely uncorrelated with firmness, fruit diameter, or fruit length. However, that study was smaller in scope, using only 30 named cultivars and 26 unselected F1 progeny.

The potential for genetic gain in fruit dimensions and fruit weight, two highly correlated measurements of fruit size, was perhaps the highest among all focus traits due to large additive and dominance effects (Table 2). These results are consistent with previous sweet cherry studies that showed high correlations between fruit size measurements and high *H*^2^ [18,78–80]. In those studies, six putative QTLs influencing fruit size in cherry were identified and together accounted for 76–88% of the phenotypic variance. Because fruit weight was highly correlated with fruit dimensions in the present study (Table 3, Fig. 4) and can be evaluated rapidly, we considered it an effective proxy for fruit dimensions and general fruit size.

The high broad-sense heritability for firmness (0.77) (Table 2) was consistent with estimates from a study conducted on a biparental population in which *H*^2^ was estimated at 0.78 to 0.85 [80]. In our study, the moderate positive correlations (r = 0.51) between fruit firmness and fruit dimensions among the unreleased progeny suggests genetic linkage among loci influencing these traits. This outcome was in contrast to that of a multi-year QTL study, in which the Pearson correlations between fruit firmness and fruit weight ranged from −0.64 to −0.67 for Regina × Lapins and −0.40 to −0.15 for Regina × Garnet F1 families [80]. Those correlations are likely due to unique linkage in Regina, Garnet and Lapins. The correlations reported here may have also been influenced by the 77 progeny derived from the three wild parents: MIM 17, MIM 23, and NY54. These individuals all had high SSC, small fruit size, and low fruit firmness in their estimated genetic values relative to the population mean (Additional File 7).

Expectations for genetic improvement in SSC were moderately positive. Narrow-sense heritability was estimated at 0.22, typical of the other focus traits in this study, where h^2^ was most often between 0.2 and 0.3 (Table 2). Broad-sense heritability of SSC (H^2^ = 0.48) was similar to that of other stone fruit: approximately 0.50 for apricot [81], 0.72 for peach [82], and 0.49 to 0.55 for apple [33]. Previous results confirm SSC had moderately negative correlation with fruit dimensions and fruit weight (−0.55 and −0.48, respectively). Our results are consistent with previous research, suggesting that SSC is directly related to photoassimilate partitioning and hence inversely correlated with fruit size [83,84]. Titratable acidity, the second most important contributor to fruit flavor after SSC, had similar variance component proportions and predictive accuracy to SSC. Major QTLs for TA have been detected on linkage groups 1, 5, and 6, explaining 99% of phenotypic variation in an F1 biparental peach population that was segregating for a large-effect locus [85]. These QTLs have not been reported in cherry. The broad-sense heritability of sweet cherry TA was lower in this study at *H^2^* = 0.60 and *h^2^* = 0.27. However, the population used in Dirlewanger et al [85] was created expressly to detect QTLs associated with TA, which might explain its very high *H^2^*.

The large *H^2^* and *h^2^* estimated for foliar powdery mildew incidence indicated excellent potential for genetic improvement, but the lack of genome-wide dominance effects was surprising (Table 2). Powdery mildew resistance in U.S. sweet cherry germplasm was first traced to a single dominant allele in the ancestor PMR-1 [86,87]. There may be evidence for other sources of powdery mildew resistance among Pacific Northwest-adapted germplasm (Zhao et al, *In Prep*). Haploblock analysis might be required to detect dominance effects, which appeared to be absorbed by the other relationship matrices. The large epistatic component (42%) determined for this trait in sweet cherry was consistent with resistance to other plant diseases such as soybean to sudden death disease (causative agent *Fusarium virguliforme*) and rice to rice blast disease (*Pyricularia oryzae*) [88–90].

### Implications for sweet cherry genetic improvement

The improvement in prediction accuracy when incorporating epistasis into the genetic model is consistent with studies on apple, Eucalyptus, wheat, cassava and maize [45,68,91–96]. Additive by additive epistasis is difficult to untangle from additive main effects due to selection, assortative mating and nongenetic covariances [44], all common facets of many breeding programs. The genomic relationship matrix for epistasis used here is considered to be an approximation since the assumption of random mating is not met [60,97]. The additive and dominance genomic relationship matrices used in this study were not necessarily orthogonal due to linkage disequilibrium between SNPs [27], and the modest correlations between the additive dominance, and epistatic values were evidence of covariance between the different genetic effects.

Epistasis has not typically been targeted for parental selection in genetic improvement programs, although it can be captured indirectly with additive effects if epistatic alleles are fixed through inbreeding or drift [68,98]. Allele fixation is challenging in predominantly heterozygous crop such as sweet cherry whose high heterozygosity is maintained by a self-incompatibility mechanism [99]. However, knowledge of allele phasing, a feature of the RosBREED sweet cherry Crop Reference Set, could enable the capture of valuable epistatic interactions through known allelic interactions for both clonal performance and breeding parent utility.

Distributions of genome-estimated breeding values from the ADI model (Additional File 7) reveals a broad base of genetic diversity and opportunity for cherry improvement. This study confirmed that the cultivar Moreau has lowest breeding values for harvest date, denoting earliness. Early Burlat and several unreleased offspring mature several days after Moreau. The highest breeding values for harvest date, designating late-season maturation, included many unreleased offspring with higher breeding values than the highest-value cultivar (Ambrunes), particularly among the families Fam35 and Fam30 that might be useful as parents. There are also many unreleased offspring with desirable breeding values for certain traits. Families Fam1 and Fam21 have high breeding values for SSC and TA. Families Fam35 and Fam16have high fruit weight and firmness breeding values, in addition to the cultivars Cowiche, Sweetheart and Selah. The breeding values reported here will enable breeders to identify valuable parents earlier in the breeding program than through phenotyping alone. Identification of parents earlier in a breeding program is a major application of genomic selection [100] and has been widely used for many crops including long-lived perennial trees [40,45,67,101–103].

Using genomic selection to skip a breeding phase has also been proposed or implemented in several crops including apple, loblolly pine, Eucalyptus and several self-pollinated and hybrid crops [29,102,104–107]. The genetic values among unreleased progeny and selections described here revealed several promising individuals with commercial potential (Additional File 7, results not shown for selections). Because sweet cherry maintains the same genetic composition and genetic potential through the breeding phases, genetic values obtained early in the breeding process will not change due to recombination. Knowing the genetic potential of an individual will help cherry breeders discard low-performing individuals and advance selections to the next phase with strong evidence. Knowledge of the genetic potential of a candidate selection may enable breeders to skip a cycle of field evaluation, thus increasing the pace of cultivar release and saving resources that can be diverted elsewhere. Given the lengthy time period for developing a sweet cherry cultivar, shortening this process may represent considerable savings.

## CONCLUSIONS

The genetic values and the improved understanding of the genetic architecture of important traits in sweet cherry obtained from this multi-year data set of a large pedigree-connected population represent a clear opportunity for genetic improvement. This application – estimating genetic variance components and genome-enabled genetic values – extended the original purpose of the RosBREED sweet cherry Crop Reference Set: QTL detection and validation. We plan to update the genetic models by incorporating new phenotypic data on the existing germplasm, adding new individuals and expanding the genome-wide SNP set for denser genome coverage. Further research is needed to validate the accuracy of the genetic predictions on an independent data set and to understand the extent of genotype-by-environment effects for the obtained breeding values and genetic values.

## DECLARATIONS

### Ethics approval and consent to participate

Not applicable.

### Consent for publication

Not applicable.

### Availability of data and material

All data used in the analyses are available at the Genome Database for Rosaceae at www.rosaceae.org [108] (persistent web link available upon article acceptance).

### Authors’ contributions

Julia Piaskowski was responsible for the bulk of the data analysis and interpretation. Craig Hardner advised on model building interpretation. Lichun Cai prepared the SNP data and Yungyang Zhao gathered phenotypic data. Amy Iezzoni and Cameron Peace contributed to study design and interpretation. All authors contributed to manuscript preparation.

## Acknowledgements

We gratefully acknowledge our funding source, RosBREED: Combining disease resistance with horticultural quality in new rosaceous cultivars, provided by the USDA-NIFA Specialty Crop Research Initiative (2014-51181-22378). We also gratefully acknowledge the contributions of an anonymous reviewer via the Peerage of Science review process, and Laura Giradeau for manuscript editing.

## ADDITIONAL FILES

Seven supplementary materials accompany this manuscript: Additional File 1 includes germplasm information on all individuals included in the study; Additional File 2 lists Spearman rank correlations by year and model and the number of observations for each year; Additional File 3 is the additive relationship matrix diagonals and off-diagonals, labeled by relationship to wild germplasm; Additional File 4 lists all variances and standard errors for all traits whose models converged; Additional File 5 lists percent variance for all components and heritability for all traits whose models converged; Additional File 6 is a biplot of the cultivars and ancestors; Additional File 7 lists breeding values, dominance values and genetic values for all individuals except selections and all traits whose models converged. An online app for exploring the breeding and genetic values presented in Additional File 7 is available at <https://jpiaskowski.shinyapps.io/cherry_gebv_xplorr/>.

## ADDITIONAL FILE CAPTIONS

**Additional File 1**: Individuals from the RosBREED sweet cherry Crop Reference set used in this study. “Self” refers to individuals derived from self-pollination, and “Unk” means that at least one parent is not known (file extension is.csv).

**Additional File 2**: Number of observations (N) and Spearman rank correlations (ρ) between genetic values derived from a single year and the multi-year genetic values using the ADI model (Panel A) or the phenotypic data (Panel B), genetic values derived from the reduced models and the full model (Panel C), and breeding values derived from the reduced models and the full model (Panel D) (file extension is.xlsx).

**Additional File 3**: Histogram of the diagonals and off-diagonal from the additive relationship matrix (file extension is.png).

**Additional File 4**: Variance component estimates and standard errors for all RosBREED sweet cherry traits (file extension is.csv).

**Additional File 5**: Variance component percentages for all RosBREED sweet cherry traits (file extension is.csv).

**Additional File 6**: Biplot of genetic values among sweet cherry cultivars and their ancestors using the correlation matrix of eight traits. Trait rotations were scaled by the first eigenvalue (file extension is.png).

**Additional File 7**: Breeding values, dominance values, epistatic values and genetic values of all individuals for all traits in the RosBREED sweet cherry Crop Reference Set (file extension is.xlsx).

